# Direct Estimate of the Specificity Constant: A Possibility or a Fluke? Pre-steady-state Substrate Concentrations and Enabling Mathematical Equations

**DOI:** 10.1101/2023.04.09.536186

**Authors:** Ikechukwu Iloh Udema

**Affiliations:** Department of Chemistry and Biochemistry, Research Division, Ude International Concepts LTD (862217), B. B. Agbor, Delta State, Nigeria

**Keywords:** *Aspergillus oryzae* alpha-amylase, specificity constant, catalytic efficiency, “burst phase-like” initial rates and corresponding concentration of substrate, Michaelis-Menten constant, second-order rate constant for the formation of enzyme-substrate complex

## Abstract

**Background:** A high-ranking scientist has recently proposed the need for direct estimation of the specificity constant rather than by calculation after a separate determination of maximum velocity and the Michaelis-Menten constant (*K*_M_).

**Objectives:** The objectives of this study are to derive novel equations for the zero-arbitrary determination of pre-steady-state (PSS) concentration of the substrate suitable for PSS assays, direct estimation of specificity constant (SC) under a PSS scenario, saturating concentration [*S*_T_] of the substrate, and instantaneous initial rate (otherwise called the “burst phase-like” rate), including its corresponding [S_T_]_0_, and to quantitatively evaluate the derived equations so as to give credence to their robustness and applicability.

**Methods:** The study was experimental and theoretical. It is supported by the Bernfeld method of enzyme assay.

**Result:** The SC values from the two newest methods for the three different concentrations of the enzyme range between 2,197.546 and 11,101.74 L/g min in one of the methods and 2,185.649 and 13,860.014 L/g min in the other method. The sub-*K*_M_ values of the SC for the three different concentrations of the enzyme range between 1304.368 and 7943 L/g min. The burst phase-like initial rate, *v*_0_, and corresponding [*S*_T_]_0_, respectively, range between 14.26 and 55.448 micro-mol./min and 0.171 and 3.752 g/L.

**Conclusion:** The derivation of the equations for the direct calculations of SC in conditions that validate the reverse and standard quasi-steady-state approximations was a possibility; the SC values are higher at lower concentrations of the enzyme. The concept of SC is very different from catalytic efficiency. The total absence of any calculation is impossible.

Graphical abstract figure for the direct estimate of the specificity constant
For the purpose of this study, the legends, SUB (S), ENZ (E), and PRD represents substrate, enzyme and product respectively. Lower number density (red) of the enzyme molecules in Figure 1 showing > number density of the product (light green) for the same concentration of the substrate (darker blue) than in Figure 2 implies that the catalytic efficiency in Figure 1 is > the illustration in Figure 2.

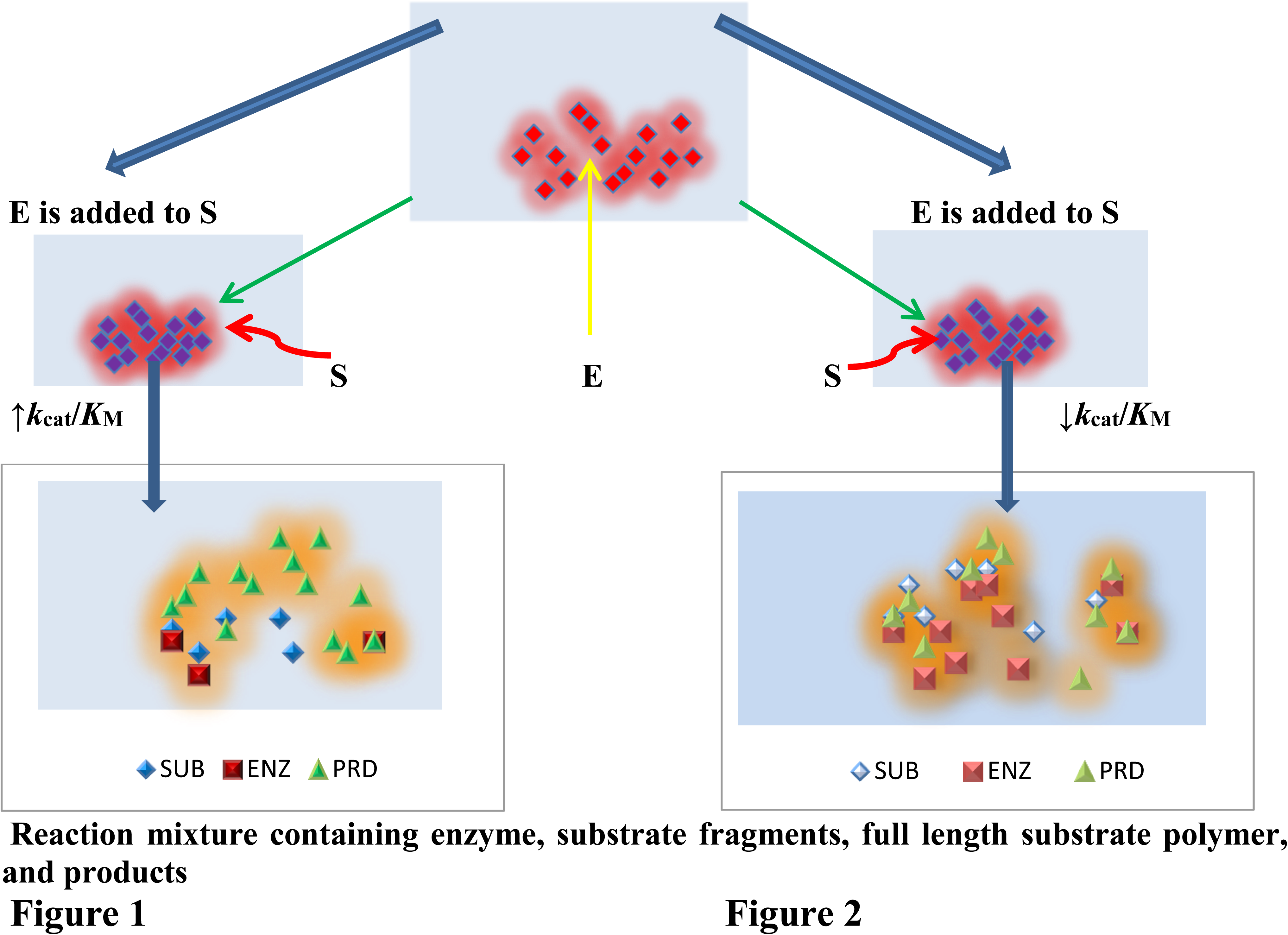

## 1.0 INTRODUCTION

Recent and past investigators have shown remarkable interest in Michaelian kinetics in the light of the need to characterize the putative enzyme being assayed either for the record or for industrial applications [1, 2]. A significant number of such investigations concern the desire to optimize the production of precursors for biofuel from the hydrolytic action of specified enzymes. Most of the time, pre-steady-state (PSS) and steady-state (SS) assays are carried out [3–7]. Burst phase, pre-steady-state, and steady-state are distinct from zero-order kinetics, and they are expected to kinetically contribute to the characterization of an enzyme whose application can be on the basis of an informed decision. These rate constants are important for proper modeling, either for further experimental or industrial design. The challenge lies in the choice of suitable substrate concentrations and the range of such concentrations. The waste to wealth concept is best achieved industrially upon application of a kinetic model; thus, research has focused on how best to convert cellulose to the precursors of biofuels [8–11] and possibly solvents for other applications in the pharmaceutical and food industries. The control of diabetes has prompted recent studies on the digestibility of recalcitrant starch and the inhibition mechanism of alpha-amylase [12–15].

Whatever is the aim of the user of enzymes, be it the saccharification and liquefaction as steps toward the production of biofuels, polar solvents, and biomass conversion before further degradation to simpler biomolecules such as trio-maltose, short oligosaccharides *etc*, and the control of simple sugar consumption in prandial scenario, there is a need to control the catalytic efficiency of the enzyme. There has been research into how best to understand the mechanisms that lead to catalytic efficiency of enzymes which is studied as reported in the literature [16–20]. The catalytic efficiency of an enzyme can be defined as its capacity to be functional at a near-optimal level in the presence of a small amount of substrate; this appears to be a function of the Michaelis-Menten (MM) constant (*K*_M_). On the other hand, there is an issue of enzyme specificity, which defines the capacity of the enzyme to selectively identify or exhibit preference for and binds consistently with the same substrate even in the presence of other substances, the substrate analogues, for instance. Here comes the issue of proficiency, which is defined in several ways, viz. A not-so-clear definition is the impression that the enzyme’s proficiency ((*k*_cat_/*K*_M_)/*k*_uncat_; where *k*_cat_ and *k*_uncat_ are the catalytic first order rate constant and equivalent rate constant without a catalyst respectively) “is the equilibrium constant for the conversion of the transition state (TS) of the uncatalyzed reaction in water and perhaps the enzyme-catalyzed reaction in water into the TSE complex [19]; the catalytic efficiency (a second-order rate constant = *k*_cat_/*K*_M_) divided by the first-order rate constant for the uncatalyzed reaction in water [17]. While catalytic proficiency stands out clearly in terms of its definition and meaning, catalytic efficiency appears to be used interchangeably with the specificity constant (SC) [18.]. It is even suggested that *k*_cat_/*K*_M_ quantifies enzyme specificity, efficiency, and proficiency [20], despite the exclusive definition of catalytic proficiency by other high-calibre scientists [17, 19].

It is important to realize that catalytic efficiency may have applications, but there have also been recent misgivings regarding the adequacy of using the ratio *V*_max_/*K*_M_ (*V*_max_ is the maximum rate of catalysis when the enzyme is saturated with substrate) as a measure of enzyme performance, particularly in the context of the use of the enzyme as an industrial catalyst. Its use can result in a misinterpretation of the performance index and can be problematic if used to select among different variants for industrial applications [21]. It needs to be noted that *V*_max_/*K*_M_ must not be allowed to be used as it is, even if the biochemist understands that a variable is missing in the simple expression given to SC: *V*_max_/*K*_M_ should be consistently stated as *k*_cat_/*K*_M_ (or *V*_max_/*K*_M_[*E*_T_], where [*E*_T_] is the molar concentration of the enzyme).

The reciprocal of *V*_max_/*K*_M_ is obtained from the double reciprocal plot and yet may not address the concern of eminent scientist [20], who prefers direct information about *k*_cat_/*K*_M_ without any form of calculation. Thus, the aim of this study is to find out whether or not a direct estimate of the specificity constant is a possibility or a fluke and, ultimately, to present pre-steady-state substrate concentrations and enabling mathematical equations. Without being prejudgmental, no one should be in doubt about the possibility of deriving a new or novel equation for the direct estimation of SC as the main objective of this study; the other objectives are to derive an equation for the zero-arbitrary determination of PSS concentration of substrate suitable for PSS assays, derive equations for SC under a PSS scenario, and an instantaneous initial rate (otherwise called the “burst phase-like” rate), including its corresponding [S_T_]_0_, and to quantitatively evaluate the derived equations so as to give credence to their robustness and applicability.

### 1.1 Significance

The study has revealed that the estimation of the specificity constant is not without any forms of calculations, though a separate estimation of the maximum velocity (*V*_max_) of the catalytic action and the Michaelis-Menten constant (*K*_M_) with the objective of finding the ratio of *k*_cat_ to *K*_M_ may not be the case, nevertheless, a single-step calculation has to be done for the determination of SC with the derived equations; the original form, the Lineweaver-Burk equation, the modified forms derived in this study and in particular the equation for the pre-steady-state case requires no more than a single-step calculation. The pre-steady-state equation replaces a not-very-accurate determination of SC in a plot of initial rate versus substrate concentrations below the *K*_M_ (the sub-*K*_M_ substrate concentrations) or in a situation where [*E*_0_] ≫ [*S*_0_], a relic of rQSSA (d[*S*_T_]/d*t* ≈ 0). The study has also given a simple equation for the calculation of what seems to be an instantaneous (or rather, a “burst phase-like”) initial rate and its corresponding concentration of the substrate. An arbitrary choice of substrate concentrations < *K*_M_ for pre-steady-state studies can be avoided by exploring the equation derived in this study.

## 2.0 THEORY

In this section, two equations are to be derived; one is based on the approach in the literature [22]. The other equation is based on a new principle. These notwithstanding, the recent direct linear method for estimating kinetic parameters are quite fortunate. In this case, no calculation is done; just the median of various points of intersection gives the direct estimate of *V*_max_ and *K*_M_. The direct linear plot [23] and the reciprocal variants [24] are graphical means of determining the kinetic parameters. The former requires separate determination of the parameters followed by calculation, whereas the reciprocal variant gives a direct value for the SC after taking the reciprocal of the *K*_M_ to *V*_max_ ratio, if initial rates were accurately generated. Thus, the reciprocal variant of the direct linear plot seems to be the first graphical approach for the determination SC. In this study, a mathematical approach and a new graphical method are to be investigated.

### 2.1 Derivation based on approach in the literature [22]

Most of the initial rate (*v*) is plotted versus the initial concentration [*S*_T_] of the substrate, but a plot of *v* versus [*E*_T_] is not out of the question. Similarly, a plot of 1/*v* versus 1/[*E*_T_] is not out of the question. Indeed, it is more appropriate to plot *v* versus [*E*_T_] as a prelude to any Michaelian investigation of any suitable enzyme. Based on the Michaelian principle, *v* is not directly proportional to [*S*_T_] because there is the presence of the latter in the denominator in addition to the *K*_M_. In this regard, Matyska L and kovář [25] strongly acknowledged that the Michaelis-Menten equation is a nonlinear equation. Hence,

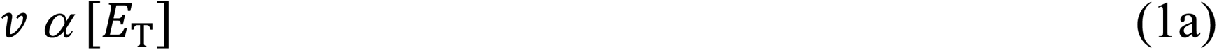

Therefore,

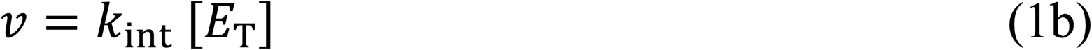

where *k*_int_ is a first order rate constant for each concentration of the substrate (S). This implies that each of the different concentrations of the enzyme up to 4 to 6 is assayed per the different concentrations of the substrate up to 6 to 10 or more. The values of *k*_int_ are different for different values of [*E*_T_]. Hence, given the following:

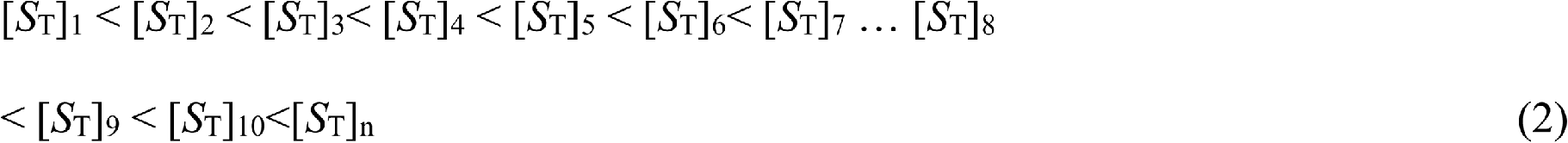

The different *k*_int_ values for different values of [*S*_T_] are:

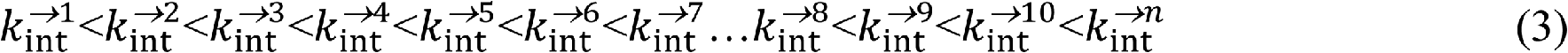

The values in Eqs (2) and (3) can be subjected to the reciprocal variant of the direct linear plot such that the median can give exactly *k*_cat_/*K*_M_. This could be very tedious and less accurate if more than six different [*S*_T_] values are used. Again, assaying for different concentrations of E (up to 4 or more) can also be tedious and take a lot of time. If the values of 1/*k*_int_ are plotted versus different values of 1/[*S*_T_], the 1/slope gives the SC value. The equation is given as:

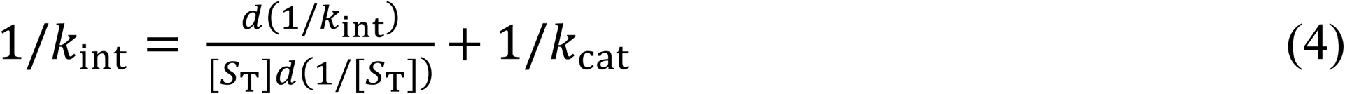

where (*d*(1/*k*_int_)/*d*(1/[*S*_*T*_]) is the slope, the equivalent of SC. The condition that satisfies standard quasi-steady-state (sQSSA) must be guaranteed for each value of [*E*_T_] in order to be accurate. A short duration (30–40 seconds) of assay is advisable in order to avoid significant substrate depletion; the concentration of the substrate at the lower part of the range may be several folds (not < 4-fold) > the highest value of [*E*_T_]. The following is derived from Eq. (4):

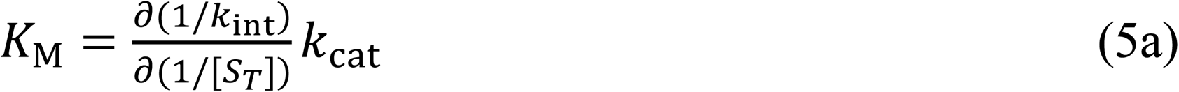

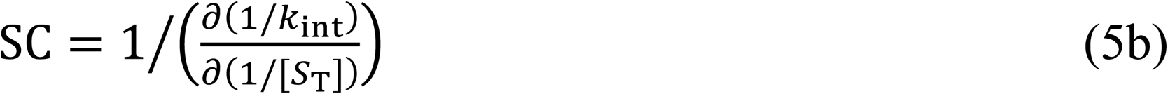

Thus, the concern for direct estimation of SC has been addressed by Eq. (5b), but the process leading to it can be tedious. Furthermore, high precision is demanded, which makes the use of automated devices inevitable. It is also very necessary to ensure that the substrate concentration regime is such that the lowest concentration of the substrate is ≫ the highest concentration of the enzyme, strictly on a mole-mole basis. A very short duration of the assay is desirable.

### 2.1 Derivation based on an alternative principle, the variation of the reciprocal of initial rates with the product of the ratios of substrate concentrations

To begin this section, the view in the literature [20] needs to be examined. Taking the equation in the literature given as

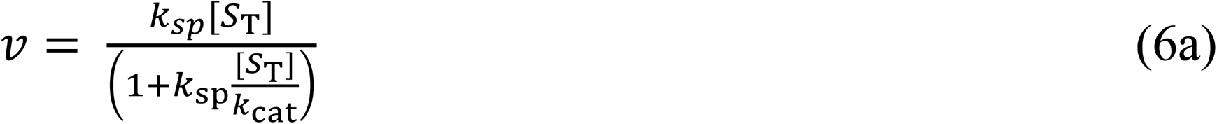

where *k*_sp_ is specificity constant. The issue with Eq. (6) is that the experimental variable *v* is not clearly defined in light of the meaning of *k*_cat_. Going by the meaning of the latter, the meaning of *v* should be a pseudo-first order constant for the formation of product at the initial stage. The comment by the author [20] that certain algebraic equations may be regarded as trivial algebra is uncalled for because the neglect of some fundamentals can lead to flawed theses even if such theses contain postdoctoral advanced mathematics that only a few can understand even in the same field at the junior level.

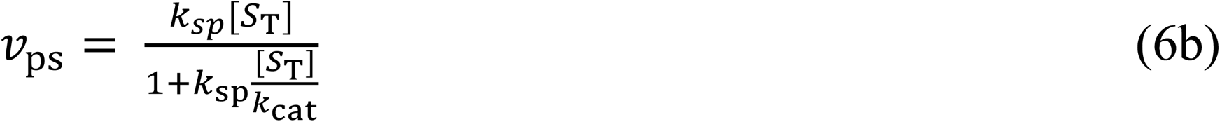

where *v*_ps_ is given as initial rate divided by the molar concentration of the enzyme.

Equation (6b) sets the standard for the subsequent derivations. But before then, a better impression regarding SC is as follows: The impression about SC is that it provides a lower limit for the second-order rate constant for substrate binding, while *k*_cat_ provides a lower limit for each first-order rate constant following substrate binding. The notion of lower limit is best interpreted in terms of the reciprocal of *k*_cat_, which gives the duration of the catalytic cycle; this, as observed in the literature [26, 27], should be equal to the sum of the reciprocals of the first-order rate constants of the individual reaction steps. This means that the individual first-order rate constant is > than the overall catalytic rate constant, referred to as the lower limit [20].

Although, the substrate concentration range may be chosen for the assay of the enzyme, there is a concentration that may be < the lowest substrate concentration of the given range. Such concentration is given as:

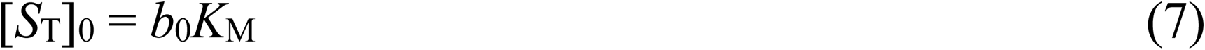

where *b*_o_ (this is < 1) is a dimensionless constant for a given substrate concentration range for a given concentration of E under a defined condition of assay, temperature, pH, *etc.* The value of [*S*_T_]_0_ is a definite substrate concentration, which may be less than *K*_M_, and which, with other concentrations less than [*S*_T_]_0_, gives initial rates that are directly proportional to the substrate concentrations. The following relationships are reasonable.

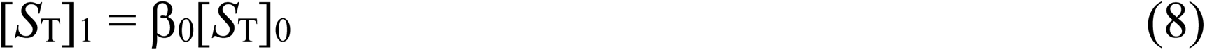

where β_o_ is the number of times [*S*_T_]_1_ is > [*S*_T_]_0_

Substituting Eq. (7) into the equation gives

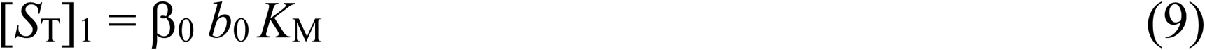

By the same line of argument, the relationship between other higher concentration of the substrate and Eq. (7) is given as follows:

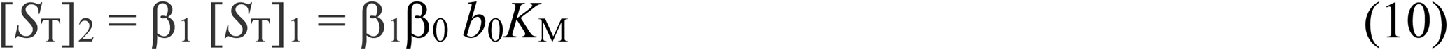

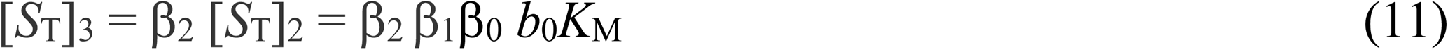

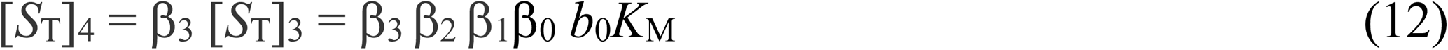

A general equation is given as:

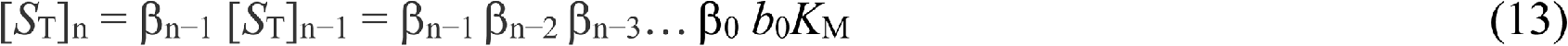

One can then write the Michaelian equation based on the above equations as follows.

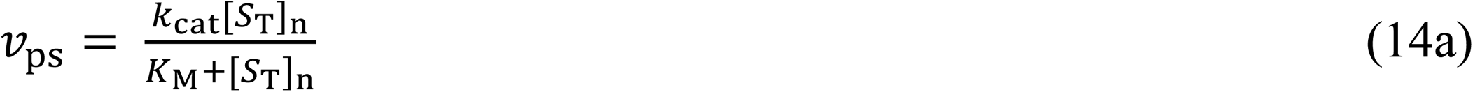

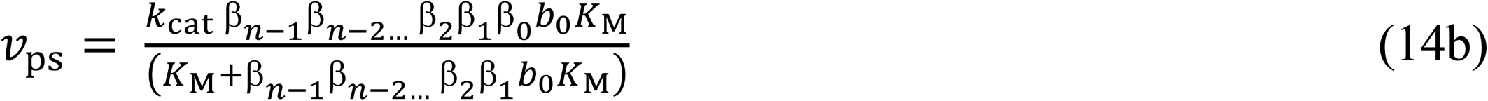

Linear transformation of Eq. (14b) gives:

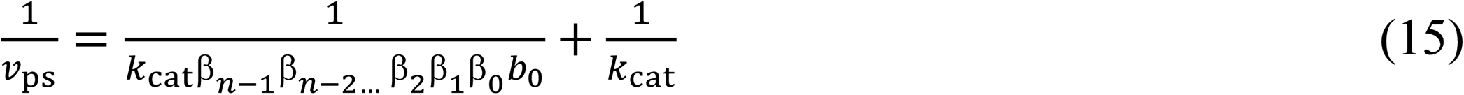

A plot of the reciprocal of *v*_ps_ versus the reciprocal of β_n−1_ β_n−2_ β_n−3_… gives intercept whose reciprocal gives the catalytic rate constant. Meanwhile, the slope (*S*_L_) from such a plot gives:

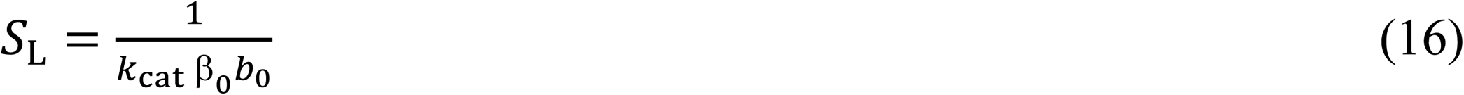

Recall that β_0_ *b*_0_ is given as [*S*_T_]_1_/*K*_M_, then Eq. (16) can be restated as:

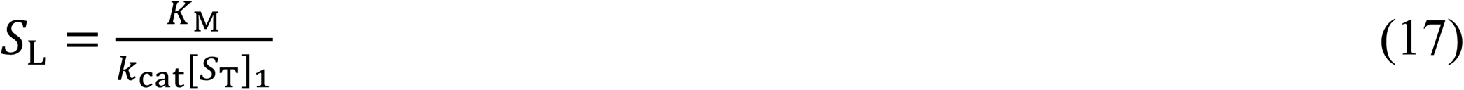

Then SC is given as:

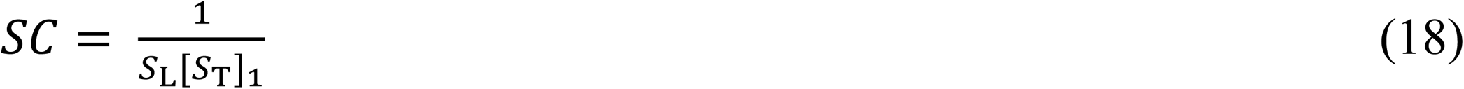

Equation (18) suggests that SC is dependent on the first and lowest substrate concentration in the range of substrate concentrations chosen.

In order to put the record straight, one should recall the premise [20] for the formulation of Eq. (15); such a premise is the assumption that there does not seem to have been an equation for the direct determination of the specificity constant despite the well-known double reciprocal transformation of the Michaelian equation by Lineweaver and Burk (LB) [28]. The challenge in the use of LB is the need to generate accurate initial rates. The plot of the reciprocal of initial rates versus the reciprocal of the substrate concentration implied in the equation [20], 1/*v* = (1/*k*_sp_[*S*_T_]) + 1/*k*_cat_, needs to make room for the following equation.

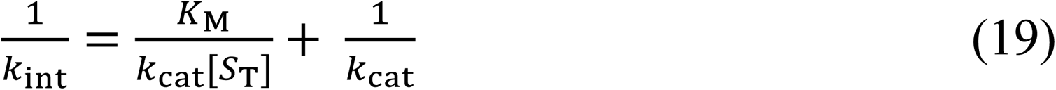

where *k*_int_ is already defined in Eq. (1b). Note that Eq. (4) is in principle similar to Eq. (19); the difference is that Eq. (4) is derived from multiple assays of different concentration of the enzyme unlike Eq. (19) which applies to a single concentration of the enzyme; both are only double reciprocal equations.

The PSS kinetic modeling requires an informed choice of a suitable substrate concentration range in addition to a suitable duration of the assay; there is a need to avoid arbitrariness in the choice of different [*S*_T_]. After choosing a suitable [*S*_T_] to [*E*_T_] ratio that should meet the requirement for a standard quasi-steady-state approximation (sQSSA), at the most basic level (this demands that the molar mass of the substrate, starch, large molecular weight protein, *etc.* must be known), the challenge, however, is that there seems to be no definite value for the molar mass (or molecular weights) of a complete starch molecule as opposed to either amylose or amylopectin. A nano-scale concentration of the enzyme with coefficients that are < 5 is preferable (for example, 1.5, 2, 2.5, 3, 3.5, 4, 4.5, and 5 nmol./L) if the molar mass of the substrate is not known with certainty. The mass-mass ratio must be avoided; otherwise, a misleading result is inevitable. In deriving the simple equations, the importance of the general equation, Eq. (13), becomes apparent in a different context. Here one plots *v*_n−1_/*v*_n_ versus [*S*_T_]_n−1_/[*S*_T_]_n_ to generate an equation of a straight line on the premise that *v*_n−1_/*v*_n_ is directly proportional to [*S*_T_]_n−1_/[*S*_T_]_n_ and partly constant to give:

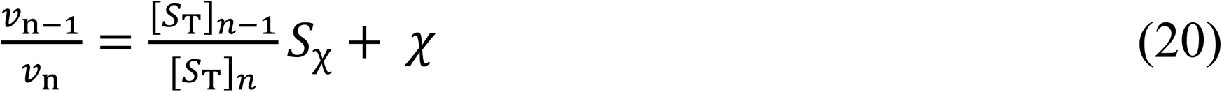

where *S*_χ_ and χ are the slope and intercept respectively. Equation (20) leads to the following:

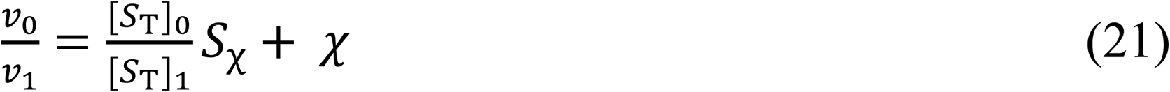

Equation (21) can be rearranged to give:

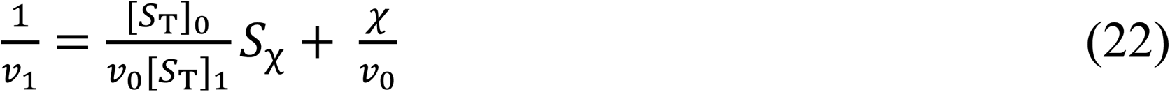

The first plot is to establish the values of *S*_χ_ and χ and since *v*_o_ and [*S*_T_]_0_ are constants for a given substrate concentration range and the concentration of the enzyme for the assay, Eq. (2) enables the determination of the initial rate before the end of the duration of the assay, which may be much less than 3 minutes: It may be a transient timescale-like the initial rate. One can rewrite a general equation for Eq. (22) to give:

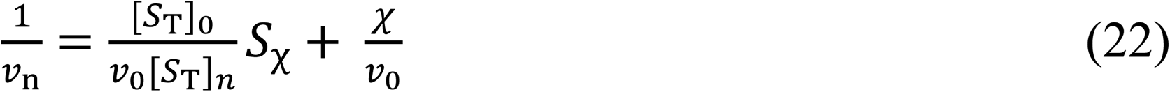

Equation (22) is simply a modification of Lineweaver-Burk [28] equation and can enable the determination of the unmeasurable initial rate just about the time the aliquot of the enzyme is added to the substrate regardless of the effect of magnetic stirring. The unmeasurable initial rate, *v*_0_ may be regarded as a “burst-like initial rate”. Given Eq. (22), the equation for *v*_0_ is derived as follows:

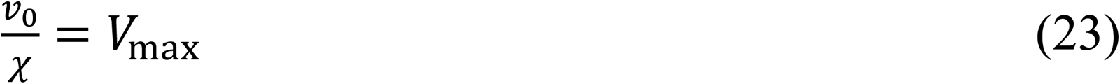

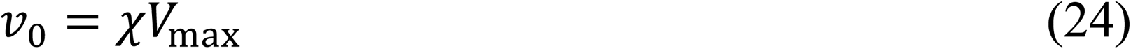

The equation for the corresponding concentration of the substrate ([*S*_T_]_0_) is derived as follows:

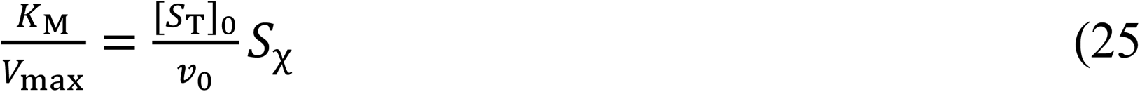

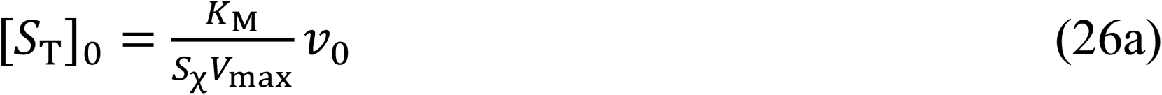

Substituting Eq. (24) into Eq. (26a) gives:

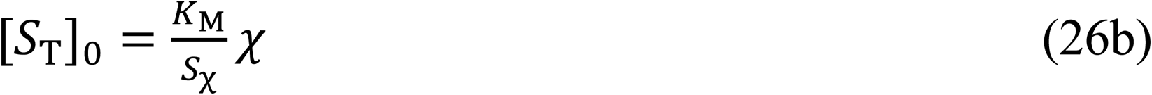

It is certain that *v*_0_ should be directly proportional to [*S*_T_]_0_ and any other values of [*S*_T_] < [*S*_T_]_0_ and ≪ *K*_M_ are also expected to observe the same direct proportionality. This is not *ad infinitum*; the best values are best defined by values < [*S*_T_]_0_. Any speculation that *v*_0_/[*S*_T_] may be ≈ SC (slightly < *V*_max_/*K*_M_) may not be out of place.

A very important lesson that should be learned from Eq. (26a) is that whenever it is correctly assumed that [*S*_T_] ≪ *K*_M_, the original Michaelian equation cannot and ought not to be rewritten as: *v* = *V*_max_ [*S*_T_]/*K*_M_. The raison *d’ être* is that such an expression is reserved for a PSS and possibly a burst phase scenario where *K*_M_ cannot be validly defined as the substrate concentration at half the maximal velocity or rate of catalysis. Instead, without insinuating any sense of novelty, the right equation should be *v* = [*S*_T_]_PSS_/*K*_d_, where [*S*_T_]_PSS_, *K*_d_, and 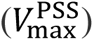 are the PSS concentration of the substrate, the enzyme-substrate (ES) dissociation constant, and the PSS maximum velocity, which is ≪ zero-order maximum velocity. Similar issues have already been elucidated in the literature [22, 26]. Recall, too, that *K*_M_ is = *K* + *K*_d_, where *K* is the Van Slyke and Cullen constants [29]. So, neither *V*_max_ [*S*_T_]/*K*_M_ nor [*S*_T_]_PSS_/*K*_d_ has any provision for *K* + *K*_d_. “ Giants in enzymology such as Van Slyke and Cullen, Michaelis and Menten need not “wake” up to address the problem or even trouble associated with the misuse of *K*_M_ often orchestrated by the living giants in biochemistry who, due to their impressive vast wealth of knowledge acquired all over the years, seem to exhibit “knowledge begins and ends with me altitude”.

Here we are again with issue arising from Eq. (26a); this is just about making *v*_0_ subject of the formula as follows:

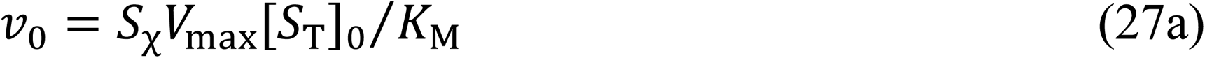

Equations (26 and 27a) are not restricted to [*S*_T_]_0_ and *v*_0_, rather every other values of [*S*_T_] that are ≪ *K*_M_ and either < [*S*_T_]_0_ or > [*S*_T_]_0_. However, d*v*_n_/d[*S*_T_]_n_ must always be = *v*_0_/[*S*_T_]_0_. How one can separately establish the value of *v*_0_/[*S*_T_]_0_ is a major research question. Therefore, with general expression (Eq. (27b)) given below one can plot *v*_n_ values versus [*S*_T_]_n_ to give a linear curve whose slope (*S*_PSSχ_) should be = *S*_χ_*V*_*max*_⁄*K*_M_. The slope *S*_PSSχ_ is simply *v*_0_/[*S*_T_]_0_. The equation is:

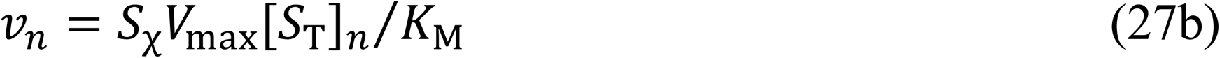

The presence of a pre-determined slope (*S*_χ_) allows the accurate estimation of *V*_max_/*K*_M_ from:

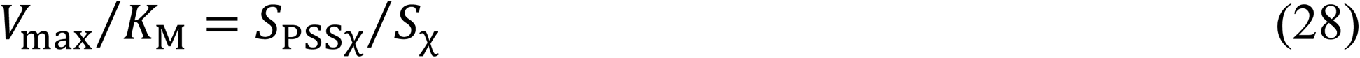

Thus, SC can be given as:

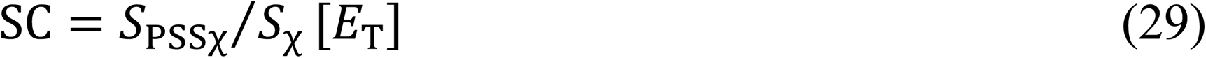

If *S*_PSSχ_is equal to *v*_0_/[*S*_T_]_0_, then Eq. (29) can be explored for the determination of zero-order SC. So far, several approaches have been derived in addition to the LW approach; none of them is without any calculation; assays have to be conducted for different concentrations of E before deriving Eq. (19); a single assay needs to be conducted if the LW approach is intended, but not without any calculation because the slope must be divided by [*E*_T_]. The most direct and minimal calculation is the reciprocal variant approach, which gives straight away 1/*V*_max_/*K*_M_ (or 1/*k*_cat_/*K*_M_ if [*E*_T_][*S*_T_]/*v* is plotted versus [*E*_T_]/*v*). Again, this is not without any calculation because one must take the reciprocal of 1/*k*_cat_/*K*_M_ in order to quantify SC. This is besides the initial calculation of *v*_ps_ (i.e., the initial rate, *v*, divided by the molar concentration of the enzyme). Equation (29) requires information about substrate concentrations that are ≪ *K*_M_. To begin with, it is necessary to point out that any substrate range much greater than [*E*_T_] will give initial rates, which when plotted versus [*S*_T_] should give a polynomial curve of the quadratic kind. Thus, given a polynomial of the kind below, a general equation for the estimation of PSS can be derived.

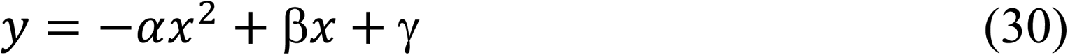

Differentiation with respect to *x* gives:

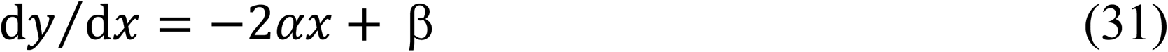

If one recalls that *v* (which is *y*) = *k*_cat_ [*ES*], then,

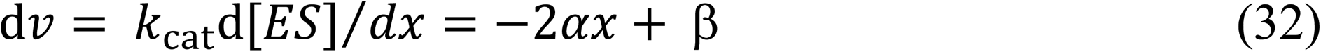

Integrating yields:

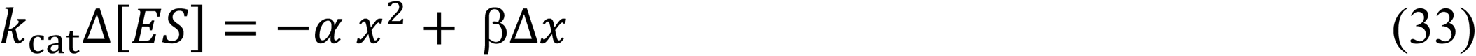

Bearing in mind that under SS condition, (Δ[*ES*]⁄Δ*t*) may be ≈ zero. Therefore,

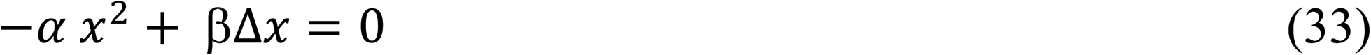

Here, *x* refers to [*S*_T_] and Δ*x* refers to Δ[*S*_T_] (= molar mass (*M*_alt_) of product × *v* × duration (*t*) of assay). The equation for PSS concentration of substrate is:

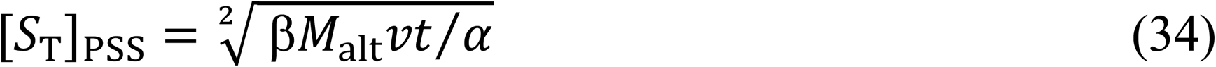

### 2.2 Another direct approach besides the reciprocal variant of direct linear plot

Another direct approach that has an attribute of generalizability is based on the alternative approach for the determination of kinetic parameters as elucidated in the literature [30]. The equation is derived as follows: The equations of maximum velocity in pre-steady-state, steady-state, and possibly post-steady-state (the zero-order scenario), and either the enzyme-substrate (ES) dissociation constant, *K*_d_ or *K*_M_ as the case may be, are:

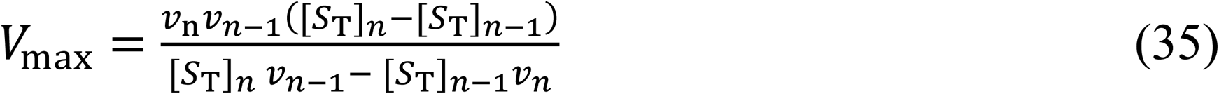

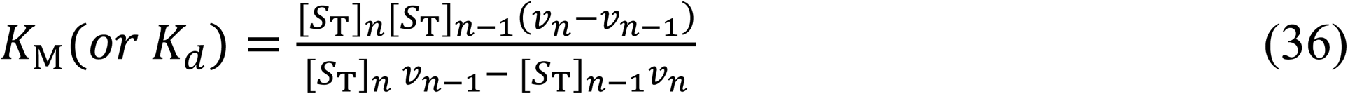

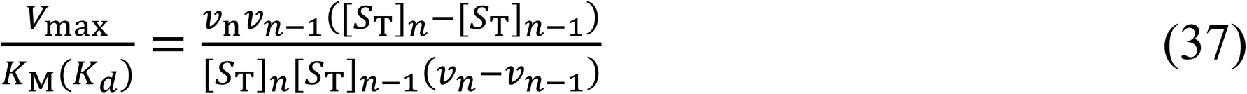

Rearrangement of Eq. (37) gives:

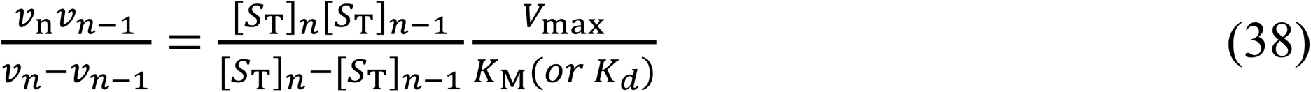

A plot of 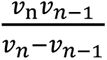 versus 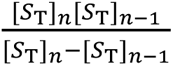 gives directly a slope equal to the specificity constant, SC. Looking at Eq. (38), one sees that preliminary calculations must be carried out, but after the plot, the slope that is read out gives the SC without further calculation except division by the molar concentration of the enzyme and multiplication by the molar mass of the substrate, if known. Note, however, that *K*_M_ must be replaced with *K*_d_ if initial rates are directly proportional to the substrate concentrations if such concentrations are < *K*_M_ and ≪ the concentration of the enzyme, as written earlier.

The alternative approach, which one can consider for a pre-steady-state scenario, is that *v*_i_ is directly proportional to the sub-Michaelian constant concentration of the substrates, and the value of the enzyme concentration is ≫ [*S*_T_]. Note once again that in such a scenario, the regression coefficient must be ≥ 0.999 and the maximum velocity should be of the pre-steady-state kind with a corresponding dissociation constant, *K*_d_; *v*_n_ could be ≅ 2*v*_n−1_ just as [*S*_T_]_n_ could be ≅ 2[*S*_T_]_n−1_. Thus,

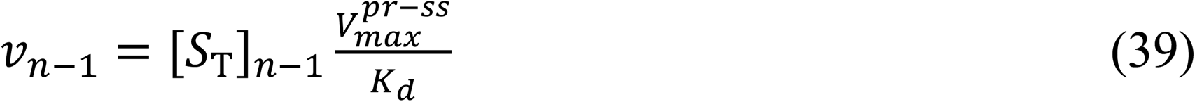

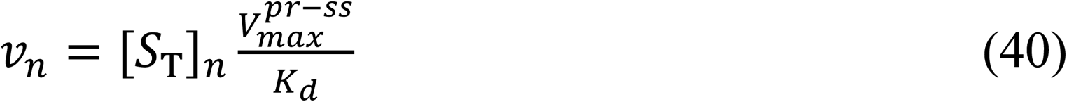

## 3. MATERIALS AND METHODS

### 3.1 Materials

#### 3.1.1 Chemicals

As in previous study [31] *Aspergillus oryzae* alpha-amylase (EC 3.2.1.1) and potato starch were purchased from Sigma-Aldrich, USA. Tris 3, 5—dinitrosalicylic acid, maltose, and sodium potassium tartrate tetrahydrate were purchased from Kem Light Laboratories in Mumbai, India. Hydrochloric acid, sodium hydroxide, and sodium chloride were purchased from BDH Chemical Ltd., Poole, England. Distilled water was purchased from the local market. The molar mass of the enzyme is ∼ 52 k Da [32, 33].

#### 3.1.2 Equipment

An electronic weighing machine was purchased from Wensar Weighing Scale Limited, and a 721/722 visible spectrophotometer was purchased from Spectrum Instruments, China; a *p*H metre was purchased from Hanna Instruments, Italy.

### 3.2 Methods

The enzyme was assayed according to the Bernfeld method [34] using gelatinized potato starch; three different concentrations of the enzyme were assayed. In this study, a mass concentration of 0.002 g/L was explored, given a mass concentration range of substrate equal to 0.3–3 g/L. The other concentrations of the enzyme that were assayed were 0.0005 and 0.0002 g/L, given a mass concentration range of substrate equal to 5–10 g/L. Reducing sugar produced upon hydrolysis of the substrate at room temperature using maltose as a standard was determined at 540 nm with an extinction coefficient equal to 181 L/mol.cm. The duration of the assay was 3 minutes. A mass concentration of 2 mg/L of *Aspergillus oryzae* alpha-amylase was prepared in Tris-HCl buffer at *p*H *=* 7. The assay was conducted at room temperature (21-23 °C.).

For reasons already elucidated in the literature [31], the equation below is suitable for the determination of the second-order rate constant if [*E*_T_] is > most, if not all, of the concentrations of the substrate that fall within the range of the substrate concentrations chosen.

The equation is however, not suitable for the case in which [*E*_T_] ≪ [*S*_T_].

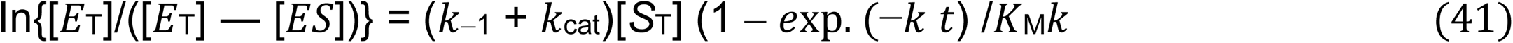

Where the pseudo-first-order rate constant, *k*, for the hydrolysis of starch is generated as described in the literature [31, 35]. The equation for the case in which [*E*_T_] ≪ [*S*_T_] is being reviewed in order to simplify the derivational procedure in the manuscript under preparation; the equation is given as:

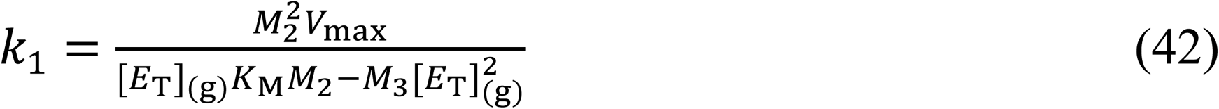

where [*E*_T_]_(g)_ and *M*_3_ are the mass concentration of the enzyme and molar mass of the substrate, the insoluble potato starch, respectively.

## 4 RESULTS AND DISCUSSION

Although the issue of inconsistency in the applicable Michaelis-Menten constant has been acknowledged and supported in a recent preprint publication [31], the starting point in this section is to reemphasize the claim that the velocity (initial rates, *v*_i_) equations of the catalytic reaction have been employed for the determination of kinetic parameters on a number of occasions outside of the conditions for which they are valid [36]. To highlight or buttress the issue, an equation was derived for a zero-arbitrary choice of substrate concentrations as explained in Eq. (34); the calculated values of [*S*_T_] based on Eq. (34) were substituted into a polynomial equation for the determination of the corresponding sub-*K*_M_ (for proper understanding, sub-*K*_M_ substrate concentrations are those concentrations that are ≪ the *K*_M_) initial rates; it must be noted that the polynomial must possess a negative coefficient of its leading term as opined in the literature [31]. However, this is strictly for illustration; otherwise, having calculated the sub-*K*_M_ concentrations of the substrate, an assay of the same concentration or a higher concentration of the enzyme needs to be conducted. A typical reverse quasi-steady-state approximation (assumption) (rQSSA) is demonstrated with a plot of the initial rates versus the corresponding sub-*K*_M_ substrate concentrations. This is illustrated by Figures 1 and 2 for the concentration of the enzyme equal to 0.0002 and 0.0005 g/L and 0.002 g/L, respectively; the rQSSA relic is that of a linear curve whose coefficient of determination is typically ≥ 0.999. This is as long as the concentration range is < than the putative *K*_M_ value of the enzyme, and better still, it should be ≪ [*E*_T_] [23, 37]. In such a scenario, the zero-order SC cannot be inferred from data points—the initial rates in particular—that either validate only rQSSA or partially validate sQSSA or by extension of the parameter domain that validates both rQSSA and sQSSA [38, 39].

**Figure 1:**
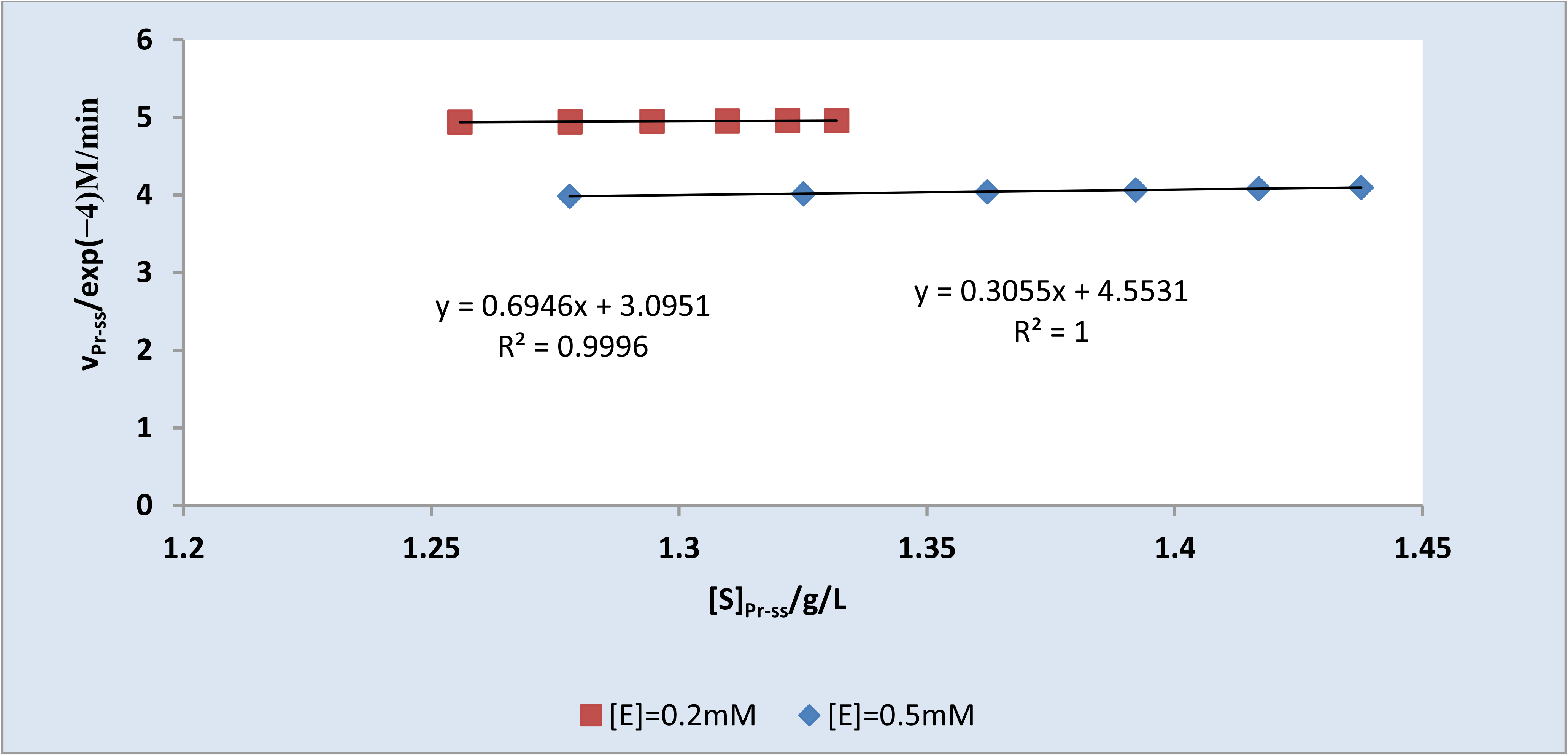
Plot illustrating the non-Michaelian characteristics of initial rates which is directly proportional to the sub-Michaelis-Menten constant concentration of the substrate where the concentrations of the enzyme are 0.0002 g/L (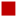) and 0.0005 g/L (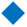). The SC for 0.0005 g/L is = 7223.84 L/g. min; SC for 0.0002 g/L is = 7943 L/g min.

**Figure 2:**
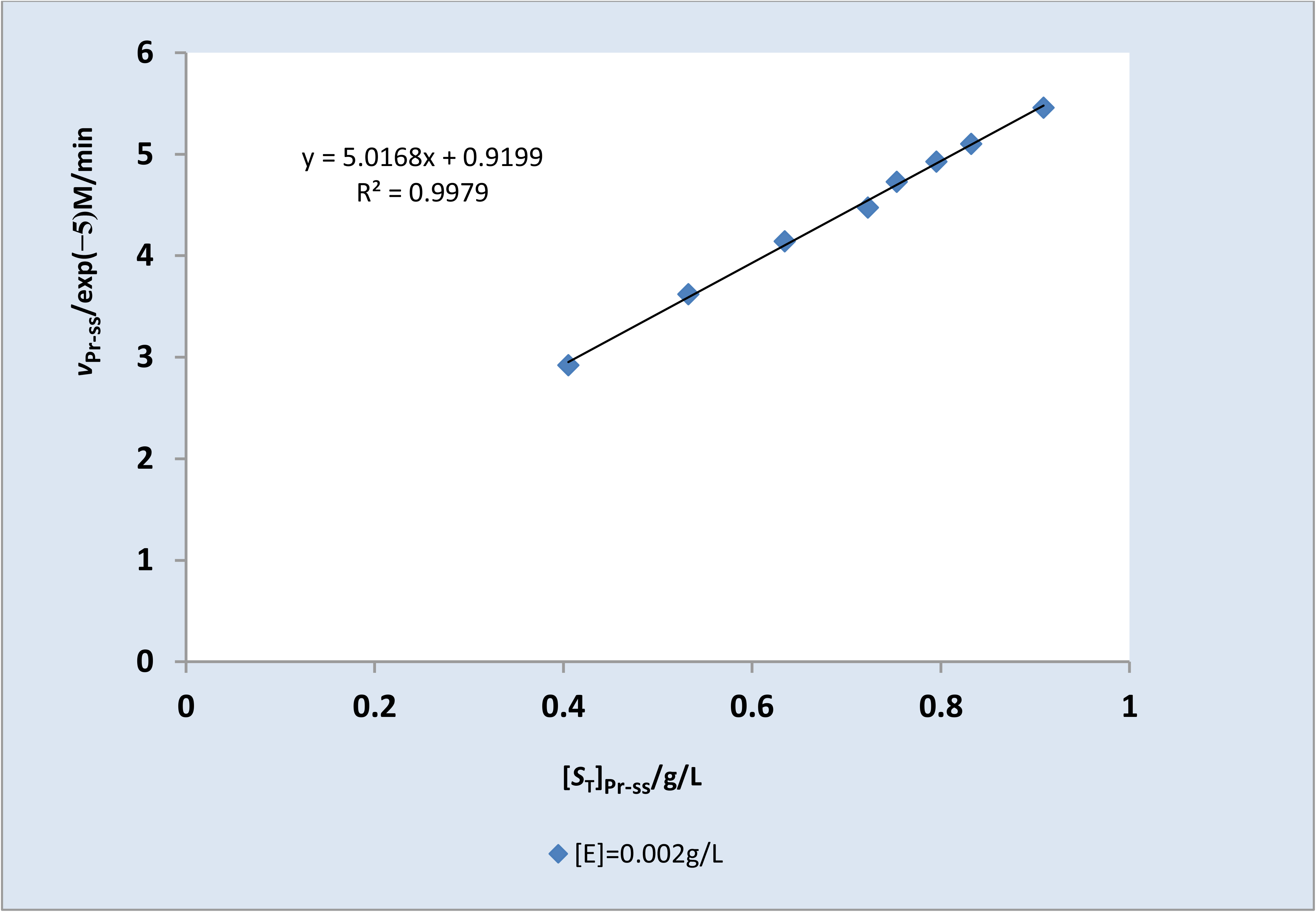
Plot illustrating the non-Michaelian characteristics of initial rates which is directly proportional to the sub-Michaelis-Menten constant concentration of the substrate where the concentration of the enzyme is 0.002 g/L. The SC value is 1304.368 L/g min.

It was hypothesized that for any given enzyme concentration that is assayed within an arbitrarily chosen substrate concentration range, as may be appropriate, there is always a “burst-like” initial rate with corresponding concentrations of substrate. These are illustrated with Figures 3 (for [*E*_T_]=0.002 g/L) and 4 (for [*E*_T_]=0.0002 and 0.0005 g/L). The “burst phase-like” initial rate and it’s corresponding [*S*_T_]_0_ are defined respectively by Eq. (24) and Eq. (26b). The latter appears to be either higher or lower than the calculated sub-*K*_M_ concentration of the substrate. With 0.0002 and 0.0005 g/L of the enzyme, the sub-*K*_M_ concentration ranges are 1.2558-1.3332 and 1.278-1.438 g/L, respectively; at those relevant values of [*E*_T_] (0.0002 and 0.0005 g/L), it is surprising to observe that the substrate concentration ([*S*_T_]_0_ = 3.752 and 2.457 g/L) corresponding to burst-like initial rates, *v*_0_ (Table 1), is > the calculated sub-*K*_M_ values. Though they are < than the substrate concentration range (5-10 g/L) chosen for the assay, they are nevertheless > than the corresponding *K*_M_ (Table 1). The situation is totally different where the concentration of the enzyme is 0.002 g/L. The [*S*_T_]_0_ value is < the Michaelis-Menten-like *K*_M._ The term Michaelis-Menten-like *K*_M_ implies that what is referred to as *K*_M_ is indeed better described as an enzyme-substrate dissociation constant applicable to a situation where [*E*_T_] is > [*S*_T_] or where [*S*_T_] is not ≫ [*E*_T_]. It is possible too, that [*S*_T_] may be ≈ [*E*_T_] [40]. Whatever the case, the burst-like initial rates for 0.0002 g/L (55.448 μM/min) and 0.0005 g/L (43.78 μM/min) were > the calculated initial rates corresponding to the sub-*K*_M_ concentration ranges, which are respectively 49.367-49.599 and 39.856-40.794 μM/min (Table 1). Again, values reported for 0.002 g/L of the enzyme were compared differently; the burst-like initial rate (14.26 μM/min) is < initial rates (29.447–54.572 μM/min), corresponding to the sub-*K*_M_ [*S*_T_] values ranging between 0.405 and 0.909 g/L. The situation might be different if the assay is carried out using the sub-*K*_M_ concentrations to generate rates rather than the calculations. The important finding is that there is a practical and theoretical guide in the determination of a lower substrate concentration regime, which may be > or < the *K*_M_ in accordance with the effect of concentration of the enzyme. The effect may imply a need to define the condition (s) that should characterize the kinetic parameters. Such characterization should be relevant to either rQSSA or sQSSA.

**Figure 3:**
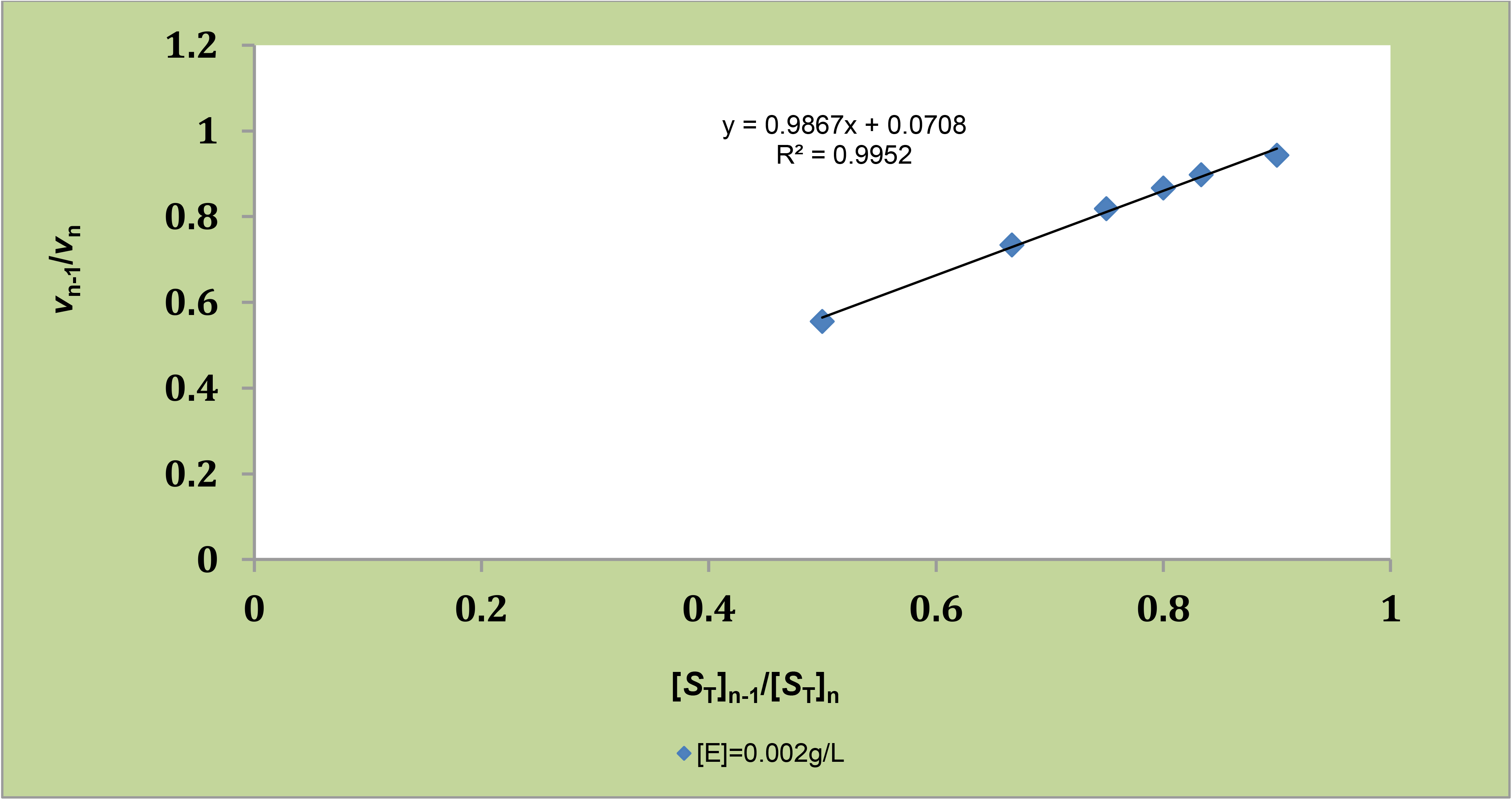
The ratio of initial rates as a function of the ratio of the corresponding concentration of the substrate, plotted versus the latter where [*E*_0_] = 0.002 g/L. [*S*_T_]_n_ > [*S*_T_]_n-1_ and *v*_n-1_< *v*_n_ where *n* is the number of assays (or “population size”) for the initial rates, *v*, and concentrations of the substrate.

**Figure 4:**
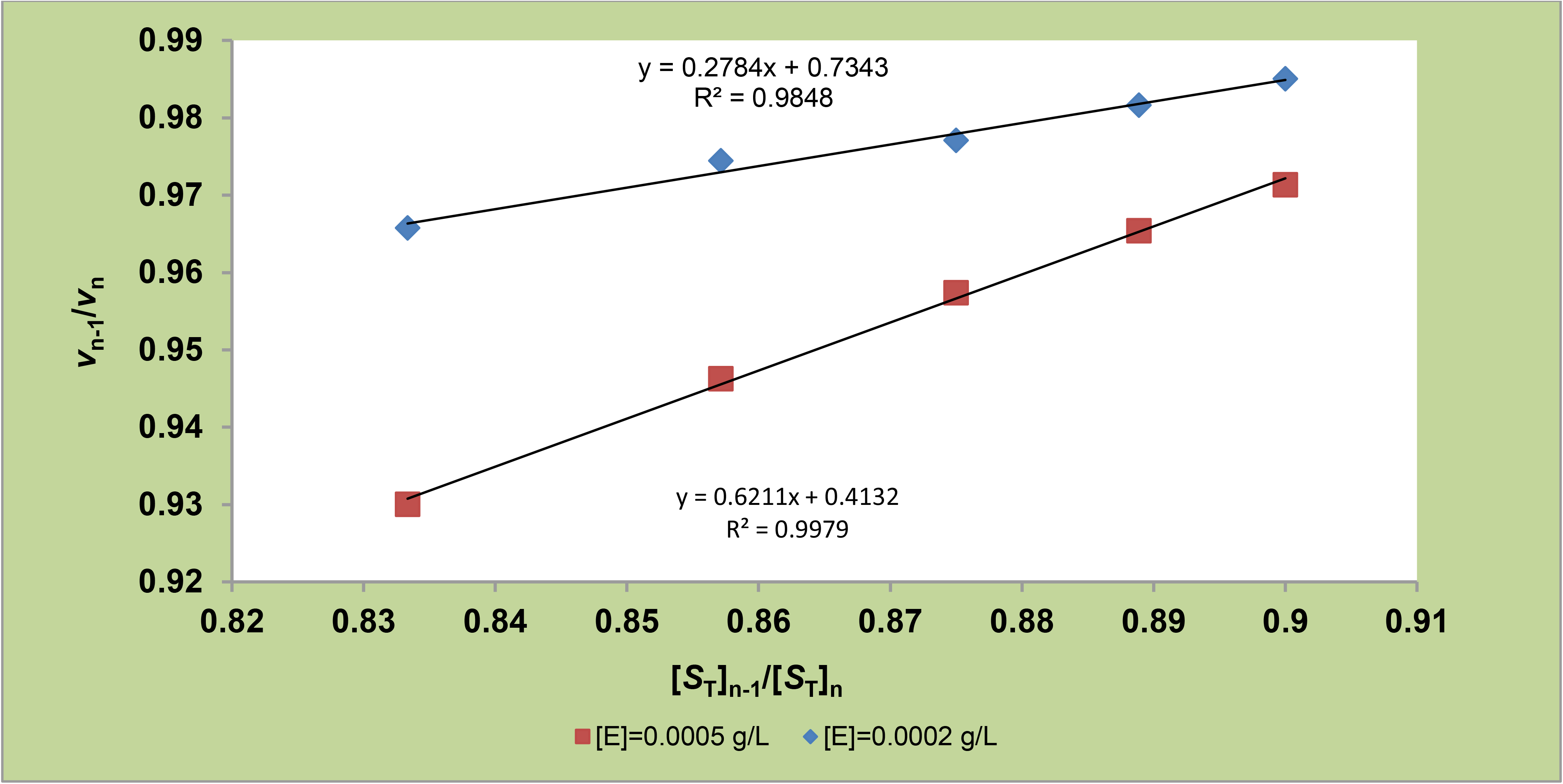
The ratio of initial rates as a function of the ratio of the corresponding concentration of the substrate, plotted versus the latter where [*E*_0_] = 0.0002 g/L (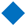) and 0.0005 g/L (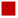).

**Table 1.** Kinetic parameters with emphasis on specificity constant.

|  |  |  |  |
| --- | --- | --- | --- |
| $[E_T]/\text{g/L}$ | 0.0002 | 0.0005 | 0.002 |
| **RVDLP** |  |  |  |
| **SC/L/g min.** | 13000 | 3687.943 | 2280.702 |
| $V_{\max}/\mu\text{M/min}$ | 78.125 | 104.167 | 166.667 |
| $K_M (K_d)/\text{g/L}$ | 1.5625 | 2.938 | 1.9 |
| **LWBP** |  |  |  |
| **SC/L/g min.** | 13800.425 | 3044.586 | 2196.688 |
| $V_{\max}/\mu\text{M/min}$ | 75.512 | 105.954 | 201.410 |
| $K_M (K_d)/\text{g/L}$ | 1.423 | 3.693 | 2.384 |
| **Equations (15 & 18)** |  |  |  |
| **SC/L/g min.** | 13860.014 | 3045.531 | 2185.649 |
| **Equation (38)** |  |  |  |
| **SC/L/g min.** | 11101.74 | 3045.12 | 2197.546 |
| $v_0/\mu\text{M/min}$ | 55.448 | 43.78 | 14.26 |
| $[S_T]_0/\text{g/L}$ | 3.752 | 2.457 | 0.171 |

RVDLP and LWBP stand for reciprocal variant of direct linear plot and Lineweaver-Burk plot respectively; $v_0$ stands for “burst initial rate”. The corresponding substrate concentration is $[S_T]_0$ .

The expression of surprises in this study outcome notwithstanding, it is imperative to understand and point out that the observed differences in [*S*_T_]_0_ and also in *K*_M_ (and *K*_d_, if applicable) values are as a result of conditions such as [*E*_T_] being at least > most of the [*S*_T_] values, approximately equal to [*S*_T_], and ≪ [*S*_T_]. The observed much lower value of Michaelis-Menten-like *K*_M_ with [*E*_T_] value = 0.002 g/L is as a result of the condition that validates rQSSA, whereby a single turnover event characterizes the catalytic cycle; this is to imply that most of the substrate molecules are subject to enzymatic action with a higher *k*_1_ value. This equally explains the view in the literature [37] that inequality cannot be held, having observed that with very high concentrations of the enzyme, “*K*_M_” is small, as in this study in which “*K*_M_” is 2.384 and 1.9 g/L (the values reported for [*E*_T_] = 0.0002 g/L) being values resulting from LWBP and RVDLP, respectively (Table 1). The values reported for [*E*_T_] = 0.0002 g/L are the lowest (Table 1).

The main objective of this study is to derive equations that are explorable for the direct estimation of SC. This is where Eq. (15) for the plot and Eq. (18) for the direct estimation of SC with a single calculation become relevant. The needed slopes are expressed in Figures 5 (for [*E*_T_] = 0.0002 and 0.0005 g/L) and Figure 6 (for [*E*_T_] = 0.002 g/L); dividing the slope by the first [*S*_T_] in the concentration range chosen gives the SC as shown in Table 1. The values of the SC compare as follows: 0.0002 > 0.0005 > 0.002 g/L. This is applicable to the *V*_max_ values. Note, however, that four versions were given. The oldest approach is the Lineweaver-Burk plot (LWBP), which gave values similar to the reciprocal variant of the direct linear plot (RVDLP) and one of the newest approaches, Eq. (18), in this research. The second and newest approach is represented by Eq. (38). To put the latter into effect, Figure 7 for [*E*_T_] = 0.0002 g/L and Figure 8 for [*E*_T_] = 0.0002 and 0.0005 g/L were created; in this case, only preliminary calculations need to be carried out. The slope has a direct value of SC. The values of SC for 0.0005 and 0.002 for all versions are similar, but not so for 0.0002 g/L, perhaps due to measurement error. In all cases, however, the SC values with sub-*K*_M_ [*S*_T_] as displayed under Figures 1 and 2 compare in the following order: 0.0002 > 0.0005 > 0.002 g/L.

**Figure 5:**
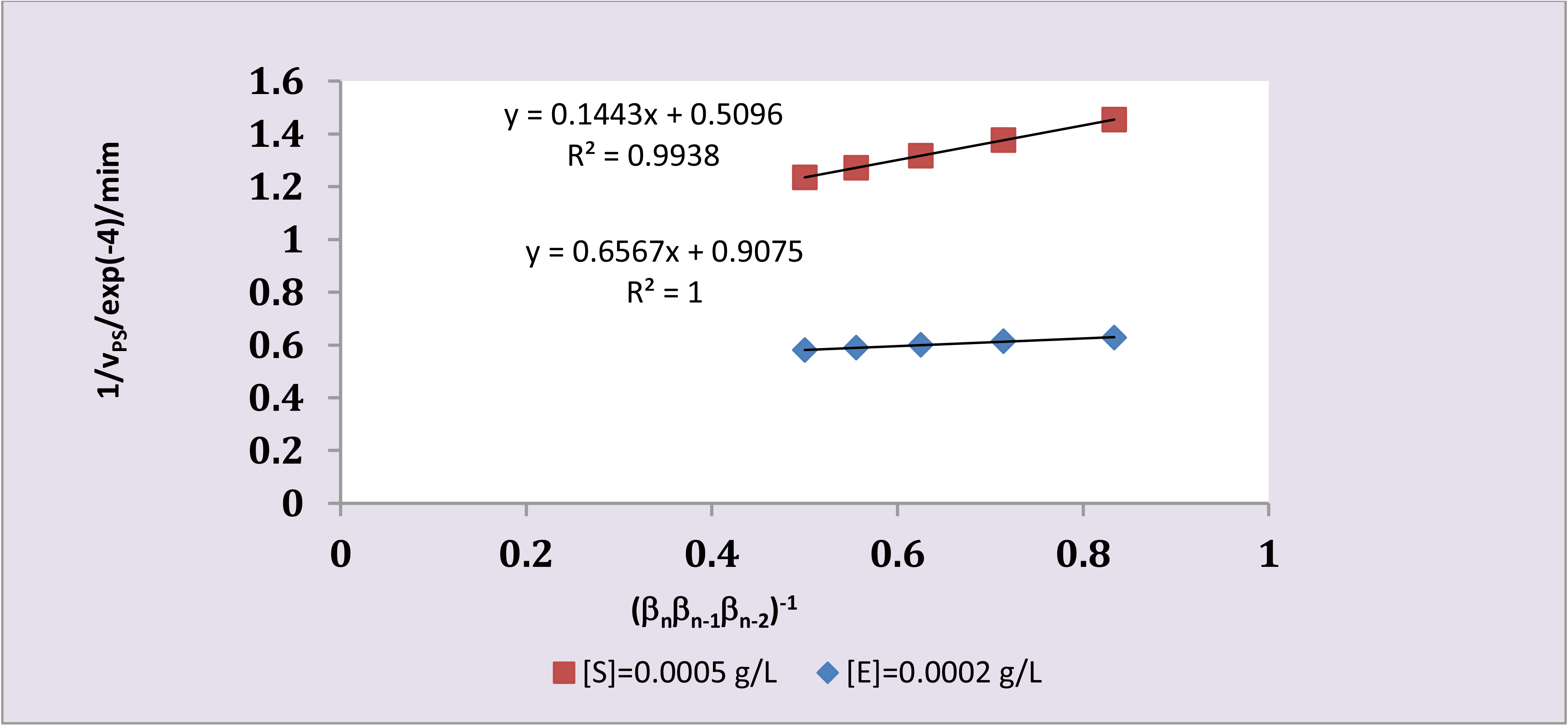
The reciprocal of initial rates as a function of the product of the ratio of substrate concentrations where the concentrations of the enzyme are 0.0005 g/L (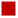) and 0.0002 g/L (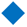). β_n−1_ = [*S*_T_]_n_/[S_n−1_

**Figure 6:**
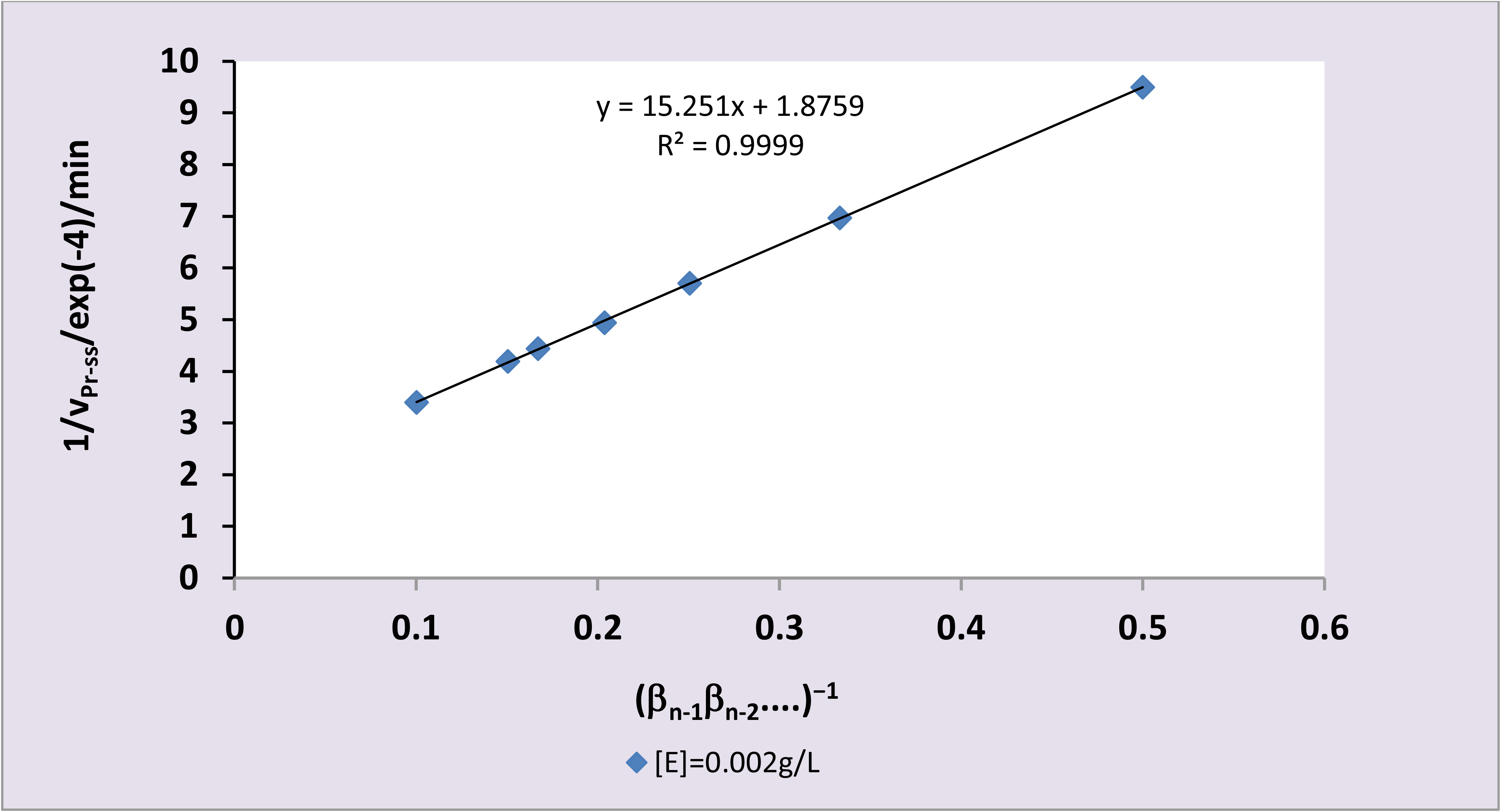
The reciprocal of initial rates as a function of the product of the ratio of substrate concentrations where the concentrations of the enzyme is 0.002 g/L. Note that taking the reciprocal of the slope (which is *k*_cat_) divide by [*S*_T_]_1_ (the first substrate concentration in the range) and multiplying by the molar concentration of the enzyme gives ≈ 8.41 exp. (−5) mol./g min.

**Figure 7:**
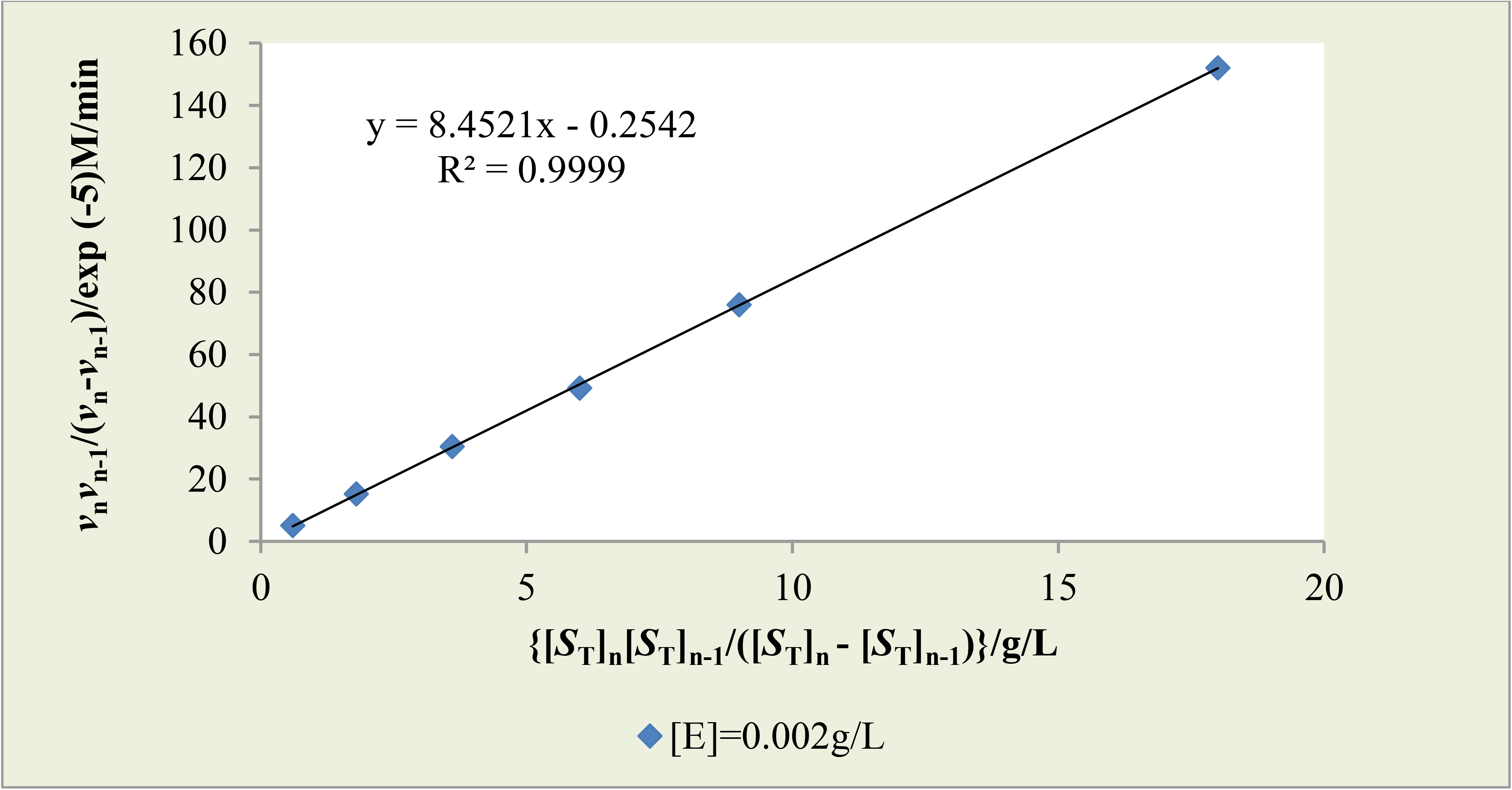
Direct determination of SC by alternative graphical approach: The ratio of the product of different initial rates divided by their difference to the corresponding product of different concentrations of substrate divided by their difference where the concentration of the enzyme is 0.002 g/L.

**Figure 8:**
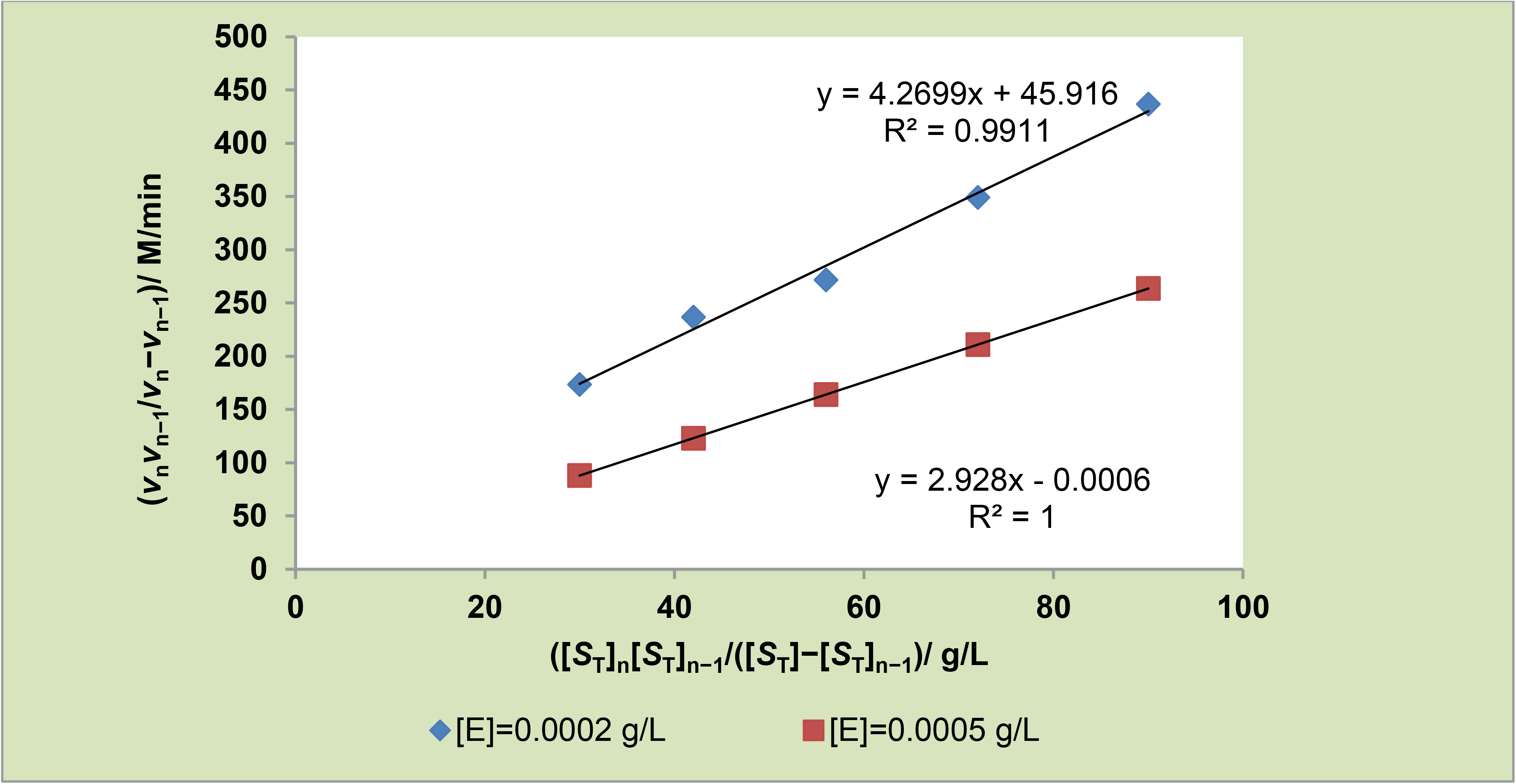
Direct determination of SC by alternative graphical approach: The ratio of the product of different initial rates divided by their difference to the corresponding product of different concentrations of substrate divided by their difference where the concentrations of the enzyme are 0.0002 g/L (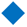) and 0.0005 g/L (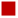).

In the construction of Figures 5 and 6, the possibility of error can be minimized or ruled out completely if care and stepwise approach is adopted in every calculation for the determination of the product of the ratio of substrate concentration. It is important that the initial rates were free of substantial error. The method is very generalizable because it can be applied in conditions that either validate rQSSA or sQSSA. The first concentration of the substrate in range of concentration chosen is very important because it forms part of the equation for the calculation of SC. Hence, it must be accurately measured.

The purpose of most industrial concerns is the optimization of production, but not without the need for any form of effective control of the production process. The products may be processed food to be packaged or drugs (prescription and over-the-counter drugs); here enzymes become extremely important in that their capacity to reduce the energy barrier (Arrhenius and Gibbs free energy of activation) can drastically influence the efficiency of the industrial processes; this is perhaps the reason why eminent scholars and research always emphasize the direct estimates of catalytic specificity. Even under normal conditions of assay (optimum conditions), some reactions may be enthalpically controlled while others are entropically controlled; the possibility that the same enzyme-catalyzed reaction is both entropically and enthalpically driven may not be ruled out. Illustrations of energy curves/diagrams abound in the literature, including Wikipedia, but this notwithstanding, Figures 9 and 10 are valuable for the elucidation of the issues in contention. When the enzyme identifies its substrate and catalytically binds to it (catalytic binding means that both substrate and enzyme are in the correct configurational and conformational orientation following effective binding), it reduces the energy barrier as the follow-up consequence; this is the specificity question.

**Figure 9:**
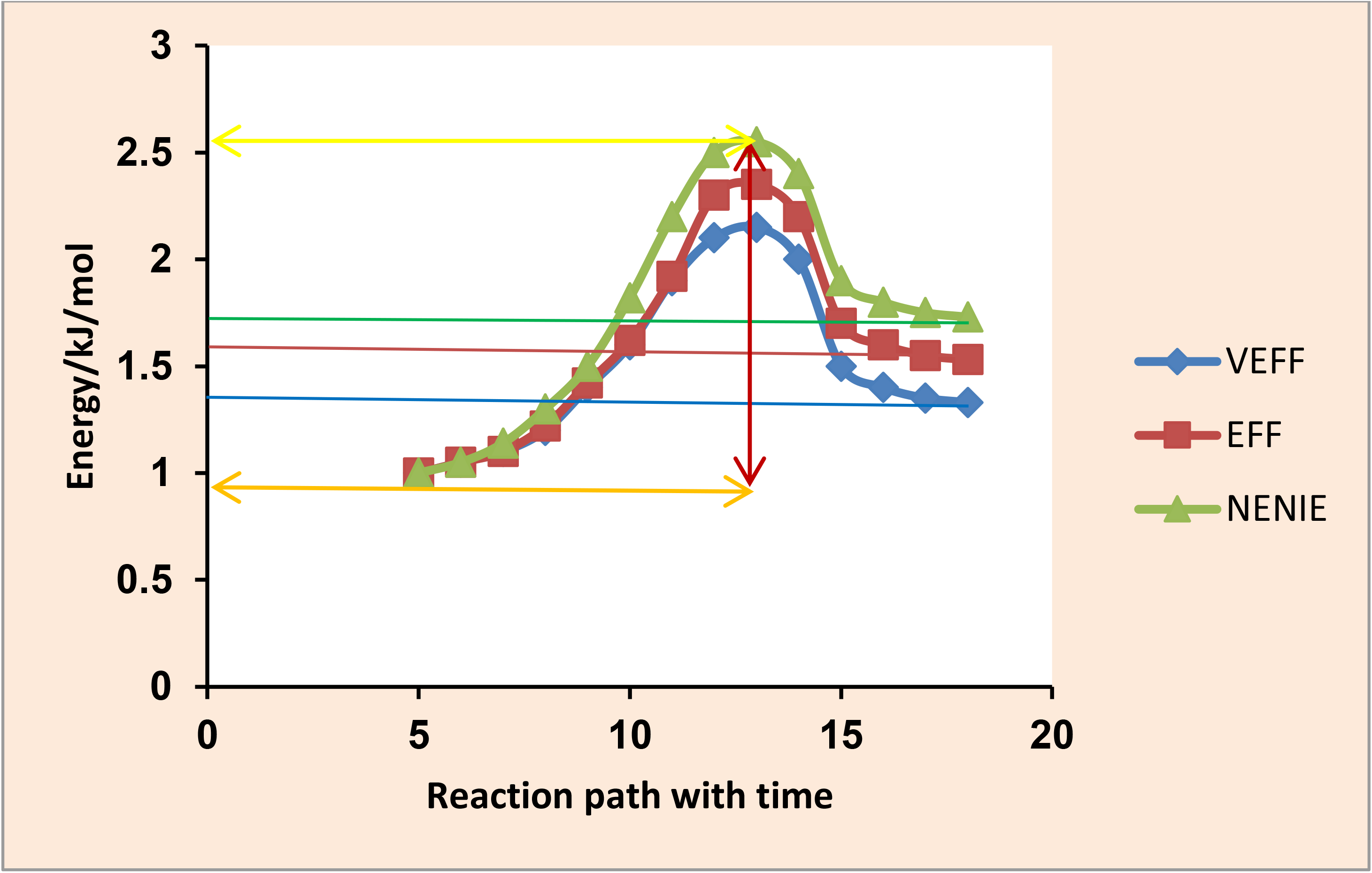
The endothermic (endogenic) case (hypothetical).

**Figure 10:**
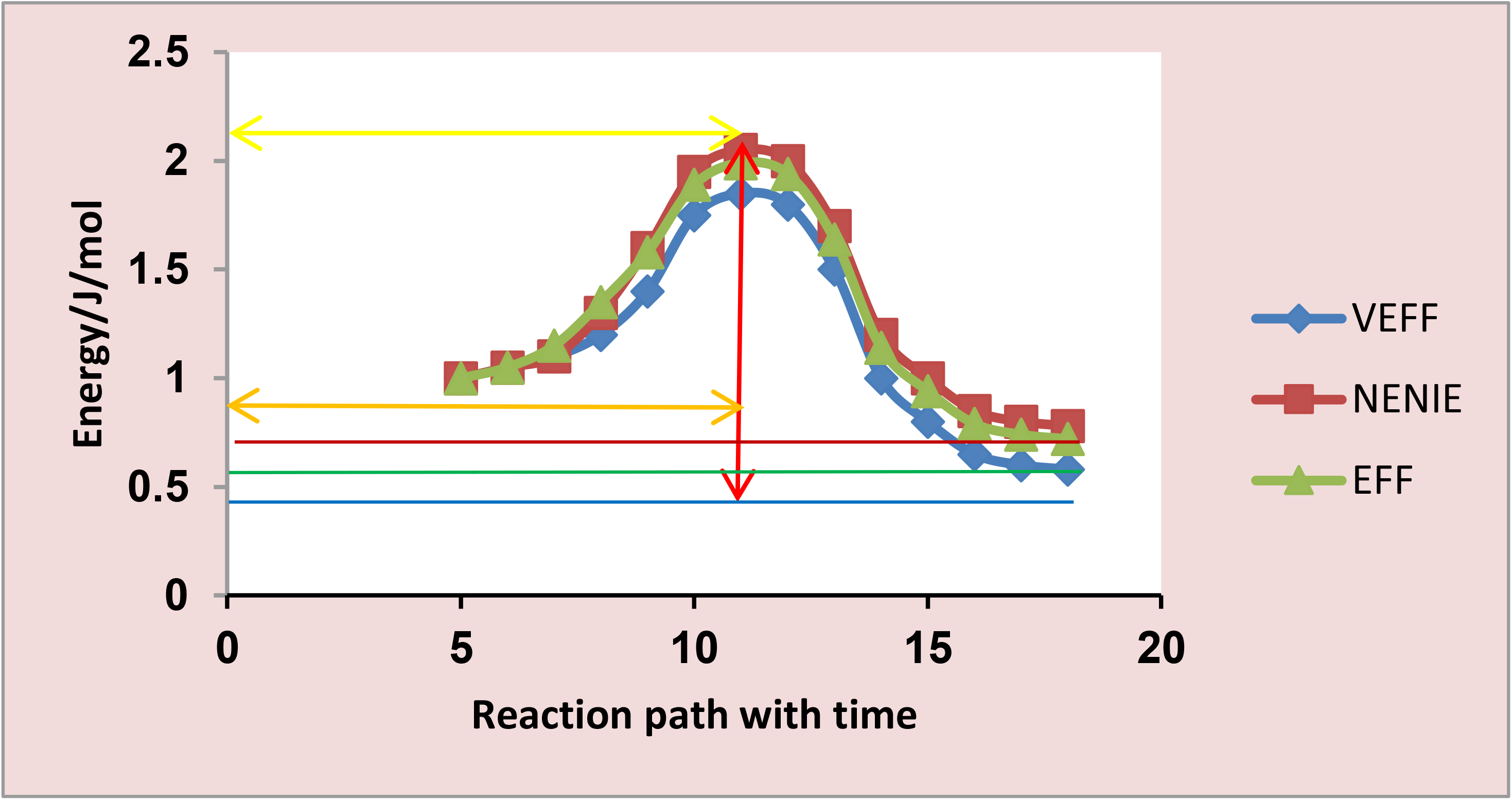
The exothermic (exothermic) case (Hypothetical).

The endothermic reactions are very peculiar with most enzyme-catalyzed reaction, notable of which is alpha-amylase amylolysis of glucans, if in particular the reaction conditions falls outside the usual. The blue curve, expresses the fact that the enzyme substantially reduced the activation, thereby becoming very efficient (VEFF), with the highest proficiency and specificity; the red curve follows next as one that is efficient (EFF), while the green curve may illustrate a situation whereby the enzyme is neither efficient nor inefficient (NENIE). The red doubled-headed arrow illustrates the energy barrier height; the orange doubled-headed arrow illustrates the upper energy barrier height-either Arrhenius activation energy or the Gibbs free energy of activation that must be added to the reactants for the reaction to proceed; the lower double-headed yellow arrow illustrates the ground-state energy level of the reactants; while the blue, green, and oxblood horizontal lines mark off the higher ground-state energy level of the products, for the very efficient, efficient and neither efficient nor inefficient enzyme respectively. In these cases the product is less stable than the reactant.

Like the endothermic case, exothermic reactions are also, peculiar with alpha-amylase amylolysis of glucans. The blue curve, expresses the fact that the enzyme substantially reduced the activation, thereby becoming very efficient (VEFF), with the highest proficiency and specificity; the green curve follows next as one that is efficient (EFF), while the red curve illustrates a situation whereby the enzyme is neither efficient nor inefficient (NENIE). The red doubled-headed arrow illustrates the energy barrier height; the orange doubled-headed arrow illustrates the upper energy barrier height-either Arrhenius activation energy or the Gibbs free energy of activation that must be added to the reactants for the reaction to proceed; the lower double-headed orange-colored arrow illustrates the ground-state energy level of the reactants; while the blue, green, and oxblood horizontal lines mark off the much lower ground-state energy level of the products, for the very efficient, efficient and neither efficient nor inefficient enzyme respectively. In these cases the product is more stable than the reactant.

To the best of the information available in the literature, apart from the proposition by a highly respected scholar in the person of Johnson [20] in the field of biochemistry that a direct estimate of SC is desirable, there does not seem to have been any attempt in the past to derive new equations as in this study, let alone quantify the values of SC. One can, however, speculate that there is always a motivation for such a wish even if there is no evidence to that effect; chemical engineers, physical chemists, *etc.* may proffer benefits or the desirability of such. However, there is no denying the fact that the two newest approaches stand out as the most effective. This is so because, where there are many data points, a direct linear plot or its reciprocal variant becomes very cumbersome and has a high potential for error if software is not applicable. There is always a high tendency for outliers with LWBP given its imprecise initial rates.

The principle of efficiency is a regular concept in both elementary and postdoctoral classical mechanics. Thus, the quality and the desirability of an enzyme for an industrial application can be evaluated on the basis of its catalytic efficiency. Without information about the reverse rate constant, one can quickly give information about the efficiency of a chosen enzyme given the ratio of SC to *k*_1_ multiplied by 100 to give the percentage efficiency of the enzyme. As shown in Table 2, lower concentrations of the enzyme under conditions that validate the Michaelian equation and sQSSA exhibit the highest value of SC regardless of the kind of direct approach; in this study, the different concentrations of the enzyme compare as follows: 0.0002 > 0.0005 > 0.002 g/L. Apart from other engineering and technical issues, the efficiency of a reactor depends on the capacity of the enzyme to reduce the energy barrier, so that with a higher catalytic rate and a lower reverse rate constant, the SC can be very high, and consequently, the catalytic efficiency given as 100 SC/*k*_1_ could also be very high. It is without doubt that one can conclude that SC is emphatically different from catalytic efficiency; unlike SC, information about the second order rate constant, *k*_1_, and SC are needed for the determination of catalytic efficiency. The latter and SC should not be interchanged, for whatever reason.

**Table 2.** Catalytic efficiency expressed as SC as a percentage of second order rate constant, *k*_1_ for the formation of enzyme-substrate complex.

| $[E_T]/\text{g/L}$ | 0.0002 | 0.0005 | 0.002 |
| --- | --- | --- | --- |
| **RVDLP** | 84.113 | 78.874 | 4.674 |
| **LWBP** | 89.292 | 65.115 | 4.501 |
| **Equations (15 & 18)** | 89.678 | 65.135 | 4.479 |
| **Equation (38)** | 71.831 | 65.126 | 4.503 |

The table of values is an out of the application of the equation: $k_1 = (k_{\text{cat}} + k_{-1})/K_M$ ; $k_1$ is a sum of two parts. Hence a fraction of it contributed by SC ( $k_{\text{cat}}/K_M$ ) multiplied by 100 gives the percentage contribution which is equivalent to catalytic efficiency, SC.

## 5. CONCLUSION

Two new equations for the direct estimation of the specificity constant (SC) were derived. This is in addition to the equations for a non-arbitrary choice of pre-steady-state (PSS) or sub-*K*_M_ concentrations of substrate, “burst phase-like” rate, and its corresponding substrate concentration, [S_T_]_0_. The SC values from the two newest methods for the three different concentrations of the enzyme range between 2,197.546 and 11,101.74 L/g min in one of the methods and 2,185.649 and 13,860.014 L/g min in the other method. The sub-*K*_M_ values of the SC for the three different concentrations of the enzyme range between 1304.368 and 7943 L/g min. The burst phase-like initial rate, *v*_0_, and corresponding [*S*_T_]_0_, respectively, range between 14.26 and 55.448 μM/min and 0.171 and 3.752 g/L; in all cases, the lowest concentration of the enzyme possesses the highest values of the parameters. The concept of SC is very different from catalytic efficiency. A future study may focus on investigating whether or not values of [*S*_T_] < [*S*_T_]_0_ with the same concentration of the enzyme can generate initial rates that, when plotted versus such values of substrate concentration, yields an original ratio equal to *v*_0_/[*S*_T_]_0_ for a possible determination of a zero-order SC at sub-*K*_M_ values.

## ACKNOWLEDGEMENT

The management of the Royal Court Yard Hotel in Agbor, Delta State, Nigeria, is deeply appreciated for the supply of electricity during the preparation of the manuscript. The language editing service of QuillBot is deeply appreciated.

---

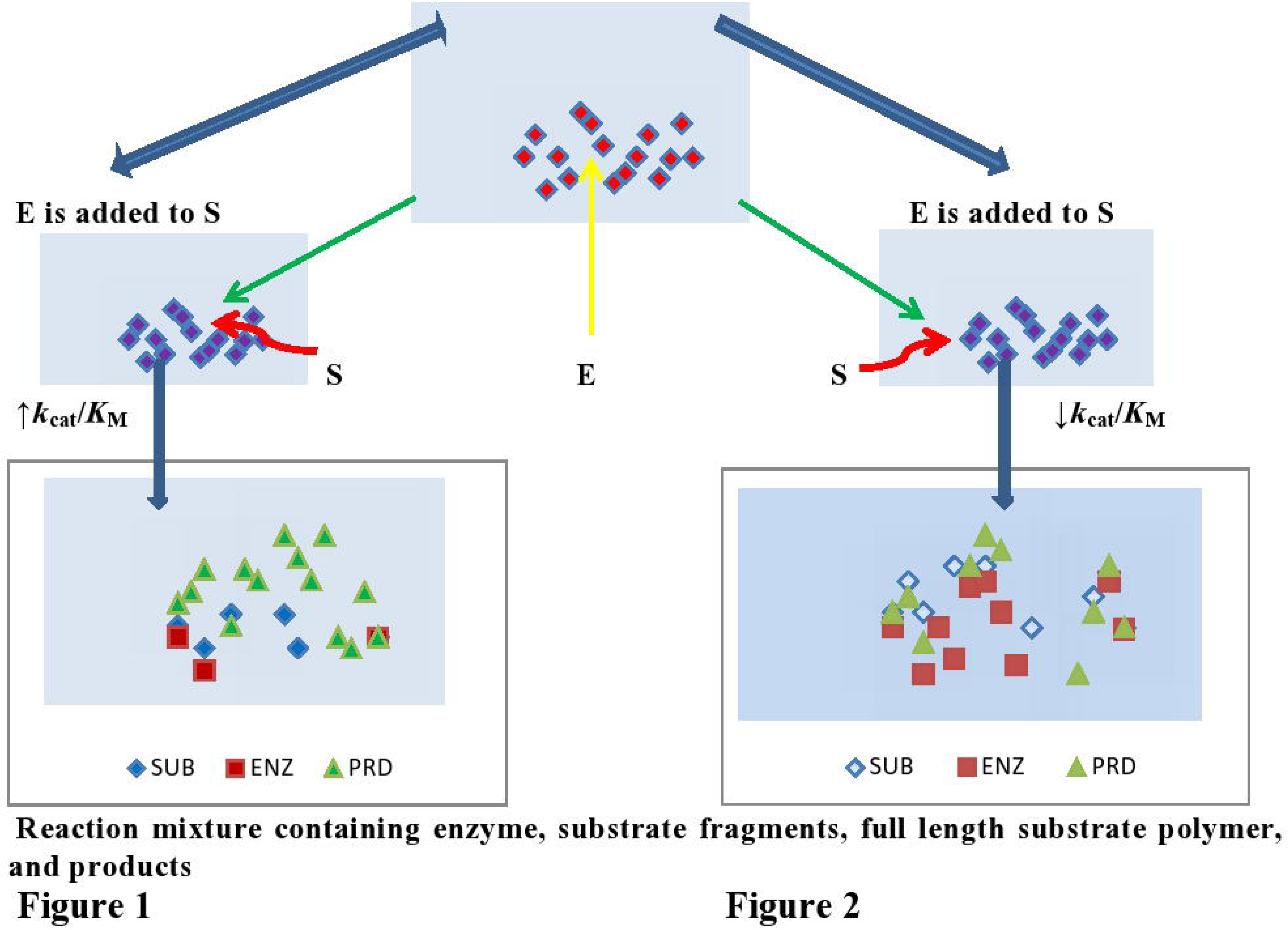
Graphical abstract figure for the direct estimate of the specificity constant. For the purpose of this study, the legends, SUB (S), ENZ (E), and PRD represents substrate, enzyme and product respectively. Lower number density (red) of the enzyme molecules in Figure 1 showing > number density of the product (light green) for the same concentration of the substrate (darker blue) than in Figure 2 implies that the catalytic efficiency in Figure 1 is > the illustration in Figure 2.

